# Creation of data-driven low-order predictive cardiac excitation models at the tissue scale in augmented state space^⋆^

**DOI:** 10.1101/2023.05.17.540314

**Authors:** Desmond Kabus, Tim De Coster, Antoine A.F. de Vries, Daniël A. Pijnappels, Hans Dierckx

**Affiliations:** Department of Mathematics, KU Leuven Campus Kortrijk (KULAK), Etienne Sabbelaan 53, 8500 Kortrijk, Belgium; Laboratory of Experimental Cardiology, Leiden University Medical Center (LUMC), Albinusdreef 2, 2333 ZA Leiden, the Netherlands

**Keywords:** data-driven modelling, surrogate modelling, machine learning, digital twin, excitable media, cardiac electrophysiology, optical voltage mapping

## Abstract

The quick and easy creation of fast in-silico models of excitable media is, for instance, needed for patient-specific predictions in diagnostics and decision making in cardiac electrophysiology. We here present a model creation pipeline that not only generates new models quickly, but also only requires data from one easily measurable spatio-temporal variable. These data may, for instance, be an optical voltage mapping recording of the electrical waves in cardiac muscle tissue controlling the heart beat. We use exponential moving averages and compute standard deviations to extract additional states from this one variable to span a sparse discretised state space. The standard deviation in a neighbourhood can be used as a proxy for the gradient. To this augmented state space, we fit a simple polynomial model to predict the evolution of this one state variable. For optical voltage mapping data of human atrial myocyte monolayers electrically stimulated by stochastic burst pacing, the data-driven model is able to describe the excitation and recovery of the system, as well as wave propagation. The data-driven model is also able to predict spiral waves only based on data from focal waves. In contrast to conventional models, with our model creation pipeline new models can be generated in a matter of hours from experiment to fitting, rather than months or years.

**Highlights:**

- Models of excitation waves can be created in minutes from experiment to fitting.
- One variable in space and time is sufficient to create a working excitation model.
- A polynomial can predict excitation waves based on useful extracted features.
- Spiral waves in heart muscle tissue can be predicted from focal wave data.

**Graphical Abstract:** 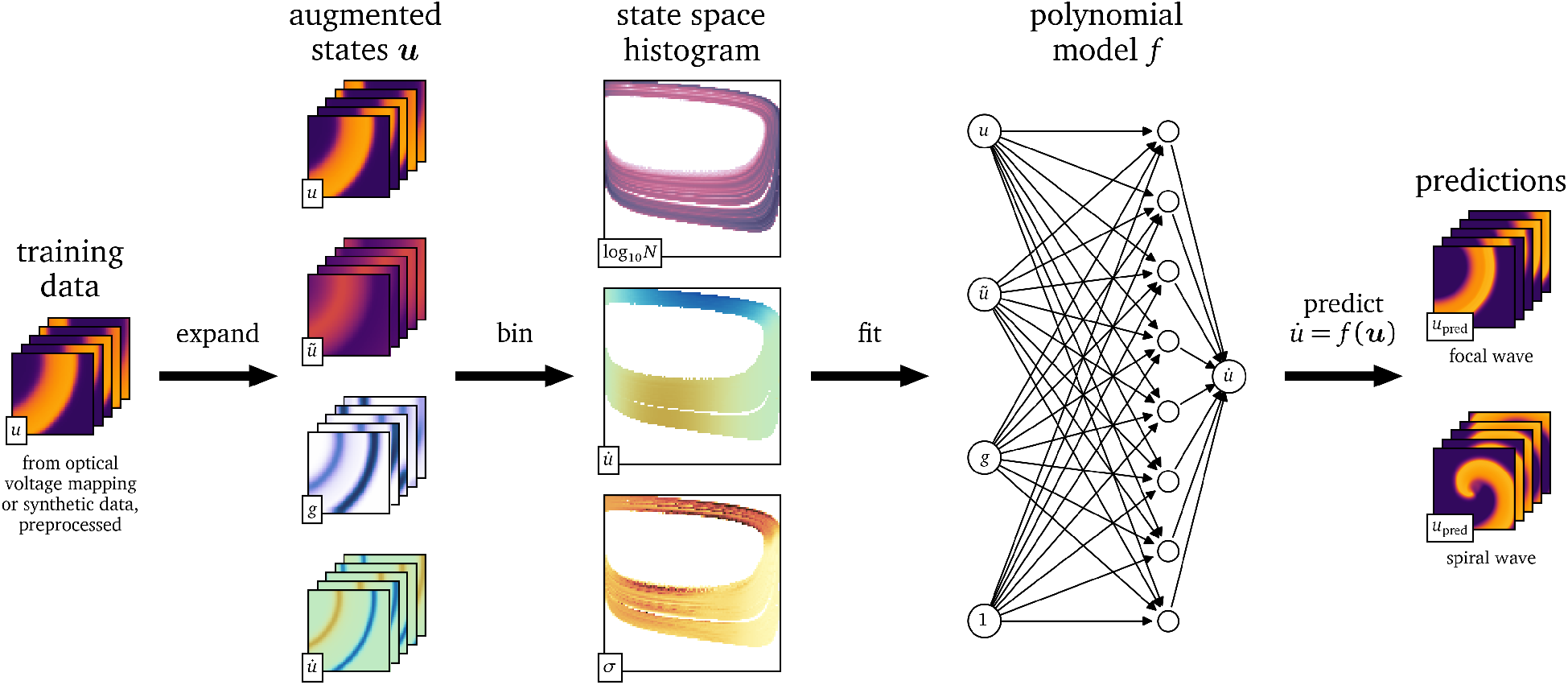

## 1. Introduction

To diagnose and treat heart rhythm disorders, a common health burden, the scientific community is creating sophisticated models for different processes in the heart with the goal of building a digital twin of each patient’s heart, i.e., a full recreation of their heart in the computer [1, 2]. One of them is the contraction of the heart which is controlled by excitation waves travelling through the myocardium [3]. An in-silico model of cardiac tissue electrophysiology is a mathematical model that is able to describe the dynamics of these excitation patterns [3].

Typically, in-silico models for cardiac electrophysiology fall into one of two categories. On the one hand, there are the mathematical models that focus mostly on the overall dynamics in the tissue [4–10]. While most of these models have continuous state variables, there are also cellular automata where each cell can only be in one of finitely many states [11]. On the other hand there are detailed models that aim to model each current relevant for the specific use case of the model across the cell membrane [12–14]. Typically, these models are fit using in-vitro data obtained from patch-clamping of single cells, or from optical voltage or Ca^2+^ mapping of monolayers or of the endocardium. Capturing cell memory is also crucial in the modelling of cardiac tissue. This may for instance be done via ionic current gates opening and closing or exponentially decaying memory variables [15, 16].

A common property in the models focussing on the overall dynamics rather than the individual ion currents is that the state variables used in them often do not have an equivalent in the physical tissue. These internal, hidden states are not fit to actual data, instead they are often chosen as a means to an end.

A so-called surrogate model is a mathematically simpler model that is fit to an existing, more detailed ionic model. For instance, Bueno-Orovio et al. [9] provide parameter sets for their model to act as a surrogate for the models by Priebe and Beuckelmann [17] and ten Tusscher et al. [18].

Surrogate modelling can be used to create patient-specific models much more quickly, such that they can be used in a clinical setting after a relatively short amount of time, e.g. during a hospital visit [19]. The current models of cardiac tissue electrophysiology are typically not fit to actual recordings of the wave dynamics. We are in this work fitting models to such data in a controlled case, i.e., dense data from isotropic monolayers, which also enables to directly compare forward simulation to experiment.

A recent step forward in in-vitro models for cardiac electrophysiology was the conditional immortalisation of human atrial myocytes (hiAMs) [20]. These well-differentiated cells address some of the shortcomings of using heart muscle cell cultures of animals for in-vitro modelling: Cells from different species have different electrophysiological behaviour, i.e., they vary in parameters such as the action potential duration (APD) and shape, the conduction velocity (CV), resulting in different wave propagation [3, 14, 20]. No in-silico version of the electrophysiological behaviour of this cell line has yet been published.

In this work, we present a general method to create data-driven in-silico tissue models from just one recorded variable over two-dimensional space and time. The method is more generally applicable and faster than the conventional creation of new models. Our method requires only minimal assumptions and works by expanding the data to a multi-dimensional state space. We both re-create an existing in-silico model with our method as a surrogate model, and use this method to create an in-silico tissue model for hiAMs only from optical voltage mapping (OVM) data. Note that our model creation pipeline is not specific to this cell line and may be applied to OVM data in general. The steps of our method are outlined graphically in Fig. 1. In section 2, we describe how the data are acquired (2.1), how they are processed, expanded with additional state variables (2.2), and how the data-driven models are finally fit to them (2.4). In section 3, we present results from the models that are generated with this method. Finally, in sections 4–5, we wrap up by summarising the method and discussing its advantages and shortcomings, as well as giving an outlook on how the models fit by our method can be improved.

**Figure 1:**
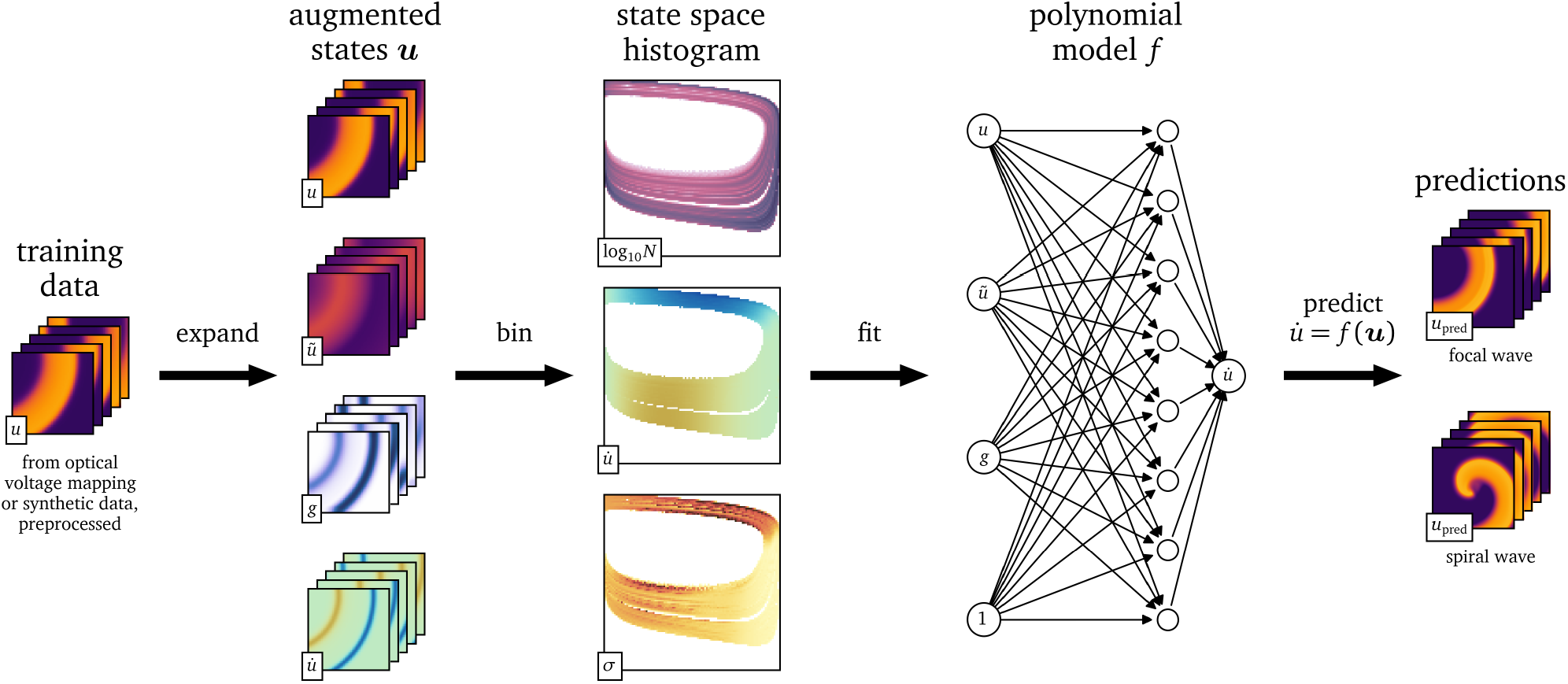
Outline of the model fitting procedure. In the first step, relevant features are extracted from the spatio-temporal data *u*. Then the state space is discretised and histogrammed. The value of the time-derivative 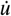 of the main variable is analysed statistically in this state space. A model can then be fit to the state space and used to predict the evolution of the medium.

## 2. Methods

### 2.1. Data acquisition

In this work, we study spatio-temporal data from two excitable media: an in-vitro data set from OVM and an in-silico data set from finite-differences simulations solving the mono-domain equations [3]. In this section, we outline how these data are generated and processed.

#### 2.1.1. Optical voltage mapping

To obtain a two-dimensional recording of the electrical activity of tissue as an in-vitro data set, we performed OVM of hiAM monolayers in 10 cm^2^-wells of six-well culture plates [20]. The recorded data consist of one variable, i.e., the measured difference in intensity of the light emitted by the voltage-sensitive dye, over space and time. Each of the 100 × 100 square pixels has a size of Δ*x* = 250 μm and the temporal resolution is Δ*t* = 1 ms between each of the 61 440 consecutive frames in time. For modelling purposes, we assume that the monolayer is approximately homogeneous and isotropic.

To excite the tissue, we place a bi-polar stimulation electrode with two poles with a diameter of 0.5 mm and inter-electrode spacing of 1 mm at the well’s boundary, and stimulate with *N*_pulse_ = 200 current pulses of random amplitude, random duration, and random timing. We call this procedure the stochastic burst pacing (SBP) protocol, inspired by stochastic pacing experiments done by Krogh-Madsen et al. [21]. The amplitudes *A*_*n*_ are sampled from a uniform distribution U (−0.8 mA, 0.8 mA). The times *T*_*n*_ = *T*_u_ + *T*_e_ between subsequent pulses are sampled from a combination of a uniform and exponential distribution

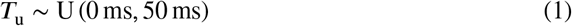

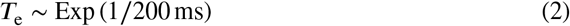

where U (*a, b*) denotes the uniform distribution on the interval [*a, b*] and Exp (*λ*) the exponential distribution with decay rate *λ*:

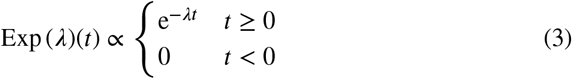

We choose this probability distribution of durations such that, on the one hand, it covers a typical range of values, but on the other, occasionally, durations are sampled that are much longer than usual.

The durations τ_*n*_ = τ_u_ + τ_e_ of stimulation are sampled from a similar probability distribution:

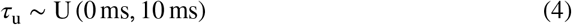

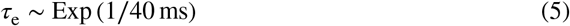

The stimulus current then takes the form:

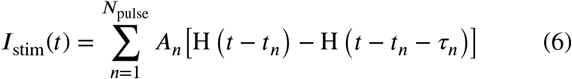

with the start time of the *n*-th pulse:

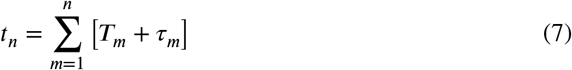

and the Heaviside function:

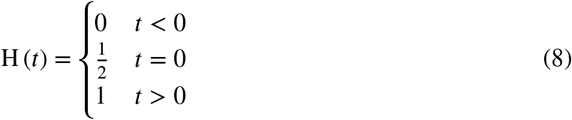

We preprocess these data in a similar way as in a previous publication [22] by Gaussian blurring with a kernel size of three grid points and rescale the data to zero-mean and unit variance.

To bring the resting state close to zero, we subtract the baseline of the data. As baseline we take the most common recorded intensity in each neighbourhood of 4 × 4 pixels over a time span of 2 s. This is based on the assumption that this most common value corresponds to the resting state.

The final preprocessing step for this data set is to smooth the data using an exponential moving average (EMA) with α = 0.1, which, for time *t* ≥ 0 with frame duration Δ*t* and the signal over time *s*(*t*) at each position in space, is iteratively defined as:

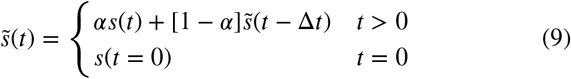

which, within first-order time discretisation, is equivalent to:

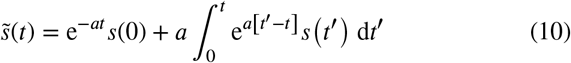

with 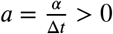.

We refer to this recording as the OVM data set and *u*(*t*, ***x***) denotes the unit-less signal after preprocessing scaled such that the resting state is close to *u* = 0 and the excited state around *u* = 1. This signal *u* corresponds to the electrical activity of the cells, i.e., it is linked to the transmembrane voltage over space and time.

Only for qualitative comparisons after a data-driven model is fit to the OVM data set, a second OVM recording is used. In this second recording, several minutes after rotor formation through burst pacing, the resulting single, stable spiral wave is observed. This experiment is also performed with hiAMs in the same well format and the same spatial resolution but a lower temporal resolution of 6 ms between frames. We preprocess these data in the same way as the main OVM data set.

#### 2.1.2. Numerical simulation

To obtain a fully controllable testing data set, we ran an in-silico experiment similar to the in-vitro experiment in section 2.1.1. We use the finite-differences method to solve the reaction-diffusion system using a rescaled version of the minimalist two-variable tissue model by Aliev and Panfilov [7], or in short the AP96 model, whose equations we will briefly summarise (eqs. 14–15).

As the diffusivity is proportional to the square of the conduction velocity (CV), we set it such that the medium has a measured CV around the CV of hiAMs [3, 20]:

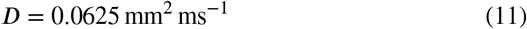

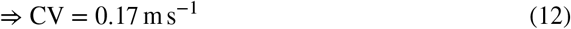

We rescale the entire model over time such that its action potential duration (APD) has a value of roughly the APD of hiAMs [20]:

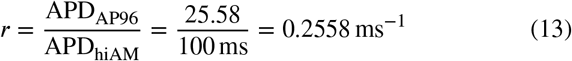

The time derivatives are rescaled by this time scaling factor *r* [7]:

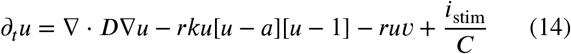

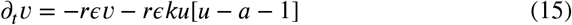

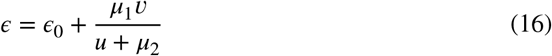

We thereby introduced the millimetre as the spatial unit and the millisecond for time into the AP96 model, which uses arbitrary space and time units. The variables *u*(*t*, ***x***) and *v*(*t*, ***x***) remain unit-less. The value *u* = 0 corresponds to the resting state and when excited, *u* approaches the value *u* = 1.

For all other parameters, *a, ϵ*_0_, *k, μ*_1_, and *μ*_2_, we use the default values, as in the original publication by Aliev and Panfilov [7]. An overview of all parameters can be found in Table 1.

**Table 1.** Parameters of the rescaled AP96 model.

| Parameter | Value |
| --- | --- |
| $a$ | 0.15 |
| $\epsilon_0$ | 0.002 |
| $k$ | 8 |
| $\mu_1$ | 0.2 |
| $\mu_2$ | 0.3 |
| $D$ | $0.0625 \text{ mm}^2 \text{ ms}^{-1}$ |
| $r$ | 0.2558 ms |

We have run a simulation over 43 104 frames at Δ*t* = 1 ms per frame at the same resolution as for the in-vitro OVM data and with the same SBP protocol. The length of the simulation was chosen such that the tissue can fully recover after the last of the 200 pulses of the SBP protocol. The electrode was placed off-centre at the side of one edge of the medium. For the time integration, we used a simple forward Euler step with numerical time step 0.1 ms and a second-order central finite-differences five-point stencil for the Laplacian ∇ ⋅ *D*∇.

The square simulated tissue is homogeneous and isotropic with no-flux boundary conditions.

We refer to this synthetic recording as the AP96 data set. We only include the variable *u*(*t*, ***x***) in the data set, as in the in-vitro data (OVM). For surrogate model fitting, we hence also only use one variable related to the transmembrane voltage of the tissue and not the internal state of the cell, which in the AP96 model is encapsulated in the second variable *v*(*t*, ***x***).

In the subsequent steps, only these data will be used, partially as training data and partially as testing data. However, we also generate auxiliary data sets for the AP96 model that are only used for qualitative comparisons with the data-driven models after fitting.

To generate a recording of a rotor using the AP96 model, we run another 2D simulation at the same resolution in space and time with initial conditions that lead to a spiral wave: The transmembrane voltage is set to a value of *u* = 0 95 within the rectangle [0, 0.5 *L*_*x*_] × [0.3 *L*_*y*_, 0.5 *L*_*y*_], and the second variable to *v* = 1 within a second rectangle [0, 0.5 *L*_*x*_]× [0.4 *L*_*y*_, 0.6 *L*_*y*_], where *L*_*x*_ = *L*_*y*_ = 100 mm. Gaussian blurring with a radius of four grid lengths is applied to avoid steep gradients at the edge of the rectangles.

We also measure restitution curves, i.e., APD and CV as functions of the CL of stimulation. For this, we run a 1D cable simulation of the model with the same spatial and temporal resolution. At the left end of the cable, we stimulate by instantaneously setting *u* to a value above the excitation threshold to trigger a wave travelling to the right. We repeatedly stimulate starting at a CL of 300 ms, decreasing in steps of 1 ms until we reach a CL where not every stimulus results in a pulse that travels through the entire cable, i.e., the CL, where 2:1 block begins.

### 2.2. State space augmentation

The multi-dimensional space spanned by all variables describing the state of the modelled medium, ***u*** = [*u, v*, …]^T^, is called state space in contrast to the physical space spanned by ***x*** and time *t*.

An in-silico tissue model usually prescribes a unique reaction ***R*** at each position ***u*** in the state space. We write a more generic version of eqs. 14–15, the mono-domain description [3]:

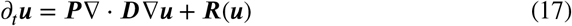

with the diffusivity matrix ***D***, where ***P*** is a diagonal matrix describing which variables in ***u*** are to be diffused.

For the two-variable AP96 model, with ***u*** = [*u, v*]^T^ and ***P*** = diag (1 0), the state space of the AP96 model is shown in Fig. 2, coloured only by the component in *u* of the reaction term ***R***. It can be seen that the second variable *v* can be used to distinguish cases with positive and negative reaction at the same value of *u*. A model that can describe both excitation from a resting state and recovery to it, is only possible with at least two variables, such that the inertial manifold can be a loop in the state space.

**Figure 2:**
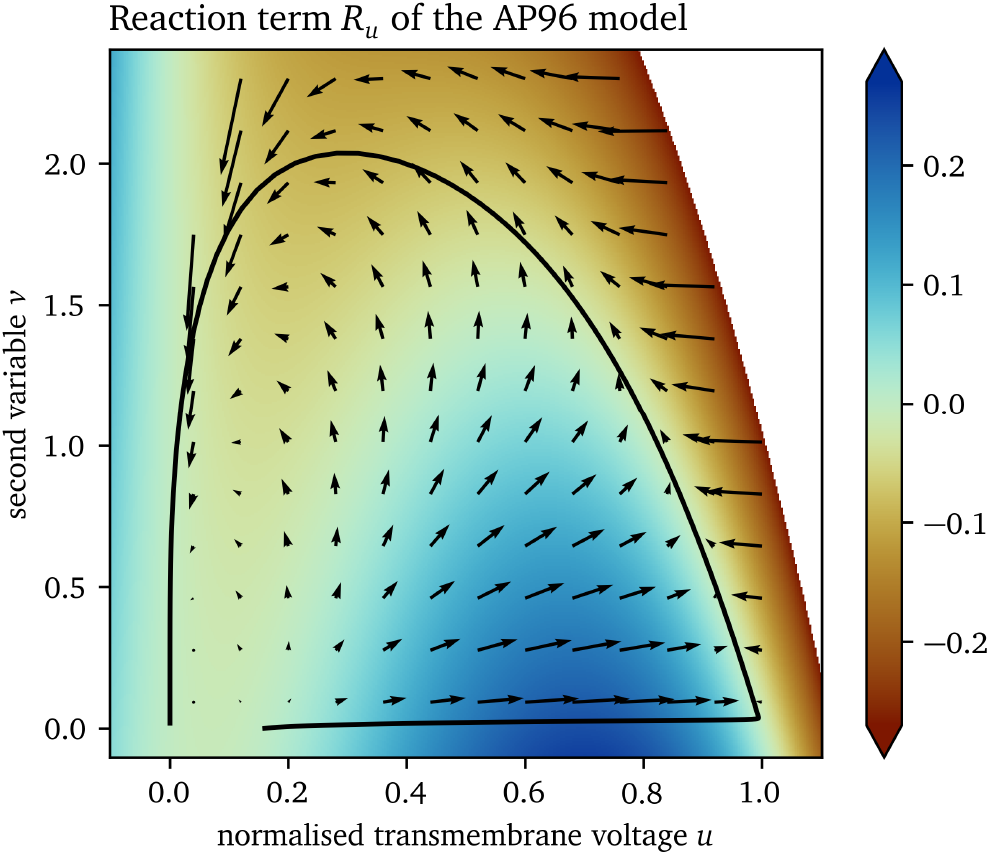
Reaction in state space of the AP96 model. The reaction term ***R*** is a vector field defining a reaction at each point ***u*** in the state space. Colouring is according to the *u*-component of the reaction term *R*_*u*_. The trajectory through the state space from excitation of one point is shown as an example.

In both types of data sets considered in this work, we only have data for one variable available, namely *u*. To still be able to differentiate between excitation and recovery, we extract additional information from *u* over space and time to use as additional states for the state space, so-called augmentations. A well-designed augmented state space unfolds the data such that cells in the tissue with different histories are identified by different augmented state vectors.

As we want to find models that can be used as forward models in a similar way as in eq. 17, we impose two more requirements on the states with which we augment the state space:

1. Augmentations should be local in space, i.e., they should only use states within a neighbourhood around Causality each point.
2. Causality must be respected, i.e., only data from previous points in time and the current point in time may be used to compute each augmentation.

The two augmentations that we found to be most useful for model creation are presented in the remainder of this section.

#### 2.2.1. Exponential moving averages

The goal of the state space augmentation is to capture that the model can evolve differently from the same value of *u*. This value may increase at the front of waves, during excitation, or decrease at the wave backs, during recovery.

One way to sense excitation and recovery is by expanding the state space with augmented states that capture the passage of time. A naïve way to do so may be to just include a time-delayed signal, which is a useful augmentation in phase mapping [23]. This is, however, not very robust to noise, as noise in the past may directly impact the present. Another option may be to use Hilbert transforms, but as their computation involves an integral over all of time, they violate causality and are therefore not meeting our requirements [24].

The EMA (eq. 9) is robust to noise and typically lags behind the signal by an amount of time inversely proportional to the constant α. It can intuitively be used to differentiate between excitation and recovery for signals *u* in-between the resting state and excited state: If an EMA with a suitable scale α is lower than *u*, it is likely that the medium is currently getting excited; if it is higher, the cells are likely recovering.

Multiple EMAs can be used to be able to capture more advanced behaviour: With an appropriately chosen α, we can differentiate between states where the tissue is activated a lot and where it is mostly resting. With such an EMA, restitution characteristics can be captured.

Here, we augment the AP96 data with a single EMA ũ with α = 3.0×10^−2^. For ũ in the OVM data, we use α = 8.0×10^−3^.

#### 2.2.2. Standard deviation as a proxy for the absolute gradient

We also conjecture that information about the slopes and higher derivatives in space of *u* is useful for a well fitting data-driven model. In the reaction-diffusion description, the Laplacian is used to capture curvature effects in space. Because it is a second-order derivative, it is very sensitive to noise. The absolute value of the gradient in combination with other state space variables can also encode much information about slopes of *u*. We use a proxy for the absolute gradient in a neighbourhood with high robustness with respect to noise. Such a proxy is the standard deviation (SD) of the signal *u* in the local neighbourhood up to a radius ***R*** around ***x***.

It can be shown analytically that the SD is proportional to the absolute gradient of a plane *h*(***x***) = ***w***^T^***x***+*b*, which evaluates to ‖∇*h*‖ =‖***w***‖. The SD in a spherical neighbourhood 𝒩 up to a radius ***R*** around the origin can be found via integration in polar coordinates *r, φ*:

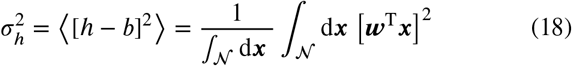

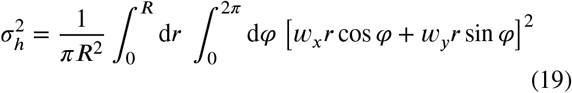

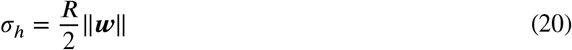

Analogously, a similar relation can be shown in 3D:

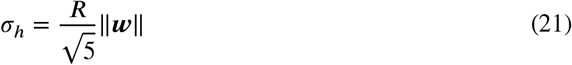

Assuming that the signal *u* varies slow enough that a linear function can approximate it well within the chosen neighbourhood, the SD can be used to compute its absolute gradient. Despite the relatively high 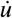 of the cardiac action potential during its upstroke, and hence ∇*u*, we can make this assumption if the resolution is sufficiently high.

Note that the SD has the useful property, that it can still be computed at or near the boundary of the medium, just based on fewer samples than for fully interior points.

To capture gradient information, we augment the AP96 data set with an SD variable for the neighbourhood up to a radius *R* = 5Δ*x* around each point ***x*** and denote it by *g*. For *g* in the OVM data set, we use a larger neighbourhood with *R* = 8Δ*x* for the SD calculation.

### 2.3. State space histograms

In the usual dynamics of the excitable systems we study in this work, not all points ***u*** in the state space are equally likely. Some state vectors ***u*** may be quite common as being part of the inertial manifold, while others might be completely absent in the data.

For example, for the AP96 model with ***u*** = [*u, v*]^T^, it can be seen in Fig. 3 that a part of the state space is not visited and most of the samples are on the inertial manifold, a closed approximately one-dimensional loop. We conjecture that in higher dimensions the state space is even more sparse, e.g. in case of detailed ionic models, some of which contain more than ten state variables.

**Figure 3:**
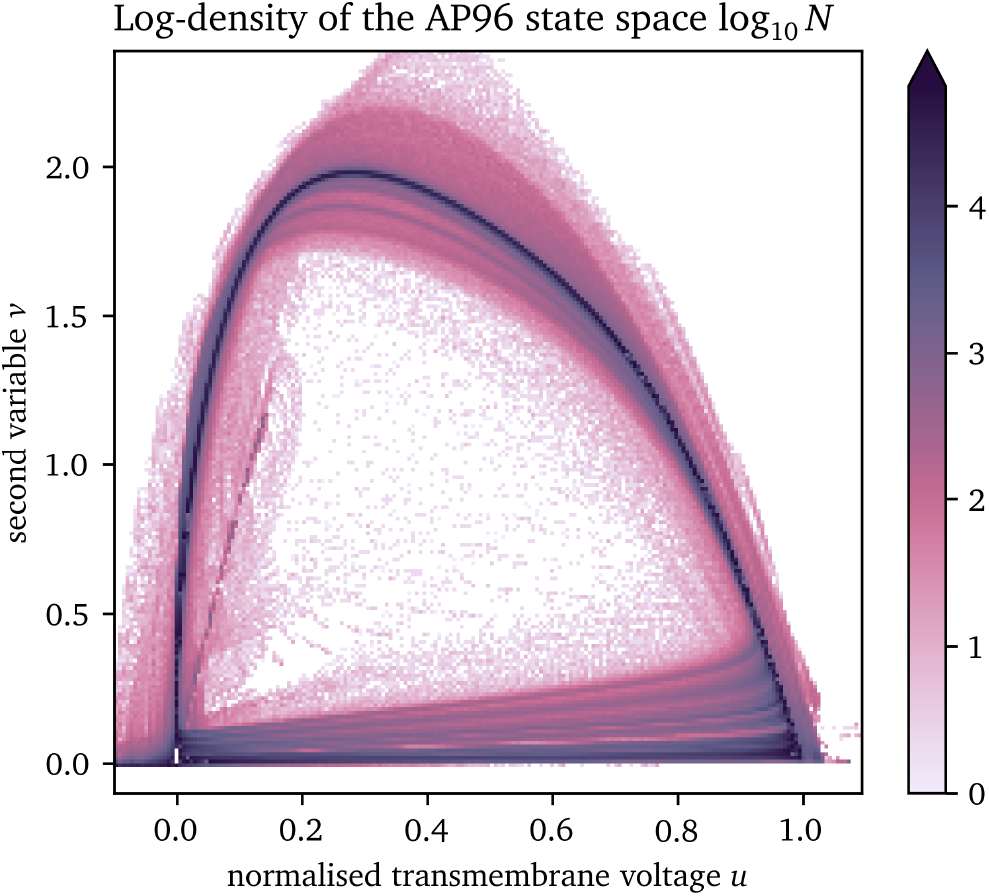
Likelihood of each point in the state space of the AP96 model for the numerical simulation generating the data set used for surrogate model fitting.

With the chosen state space expansion (section 2.2), the state vector is ***u*** = [*u, ũ, g*]^T^. We discretise the state space to 100 bins in each dimension and collect the data in a multi-dimensional histogram. This results in three-dimensional state space histograms for the two data sets with 10^6^ bins each. For efficient memory usage, these histograms are computationally implemented using multi-dimensional sparse arrays. We can then fit the models to the average value inside each bin instead of to all data points. This efficient memory management enables building even higher-dimensional state spaces, though in this work we require only three variables for now. State vectors which are completely absent in the data do not take up memory this way.

The time-derivative 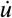 fully defines the reaction of the data-driven models developed in this work. The reason for this is that all other variables in the state vector ***u*** can be derived from the one variable ***u*** over space and time.

In a well chosen state space, for each state vector ***u***, the reaction 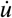 is the same, i.e., the states are well separated. Therefore, we calculate the mean value of 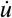 computed using finite-differences at each bin in the discretised state space, as well as its SD. This mean value of 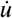 takes a similar role as *R*_*u*_ for the AP96 model and “colours” the state space in a comparable way to *R*_*u*_ in Fig. 2.

### 2.4. Model fitting

A data-driven model will be fit to the data in state space as a scalar-valued function *f* :

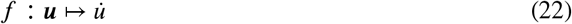

such that the model consists of the state space expansion ***u*** using eq. 9 to calculate the EMA and calculation of the SD within the neighbourhoods around each point (section 2.2.2), as well as the forward Euler step:

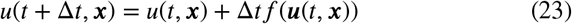

To find the optimal model function *f*, we use the least-squares algorithm to find the polynomial of a given order which minimises the loss:

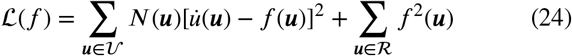

where 𝒱 denotes the centres of all bins in the discretised state space containing at least one sample, i.e., one data point, and *N*(***u***) is the number of samples in the bin ***u***. To regularise the model space in favour of more physically realistic models, we add additional co-location points *ℛ* outside of *𝒱* at which we assume *f* should be close to zero. We choose these co-location points *ℛ* by Latin hypercube sampling and only keep points for which no data is contained in the data set *𝒱* [25]. We choose the number *N*_*ℛ*_ of co-location points *ℛ* based on the number *N*_*𝒱*_ of non-empty bins in the data set *𝒱* :

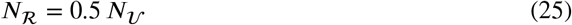

As the model function *f*, we choose to fit a low-order polynomial which is linear with respect to each of the components:

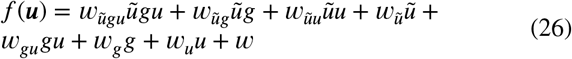

To prevent the fit model from leaving the valid state space, after each forward Euler step, we clip the predicted *u* to the range for which we have training data, the so-called valid range. Note that this also limits the EMA *ũ* to that range, while the SD *g* can still leave the range observed in the data, i.e., for larger than typical slopes.

As training data spanning the state space *𝒱* that the model is fit to, we use all frames up to just before the last pulse of the SBP in each recording. The frames of the last pulse each is then used as testing data, i.e., as a reference to compare with the predicted frames of the data-driven model. We only include points at least 20 grid points away from the boundary to minimise boundary effects on the used data.

Besides this first simulation that can be directly and quantitatively compared to the reference for both data sets, we also run simulations using the data-driven models to compare them qualitatively with the reference model, i.e., the AP96 model for the synthetic data set, and the in-vitro model for the OVM data. As the second simulation, for both data-driven models, we run a similar rectangle-based protocol to the one used to stimulate spirals for the reference AP96 model (section 2.1.2). Here, we use the same stimulation values for *u*, but *ũ* = 0.6 inside the second rectangle and *ũ* = 0.1 outside of it. Thirdly, we also run a 1D cable simulation to obtain APD and CV restitution curves for the fit surrogate AP96 model with the same restitution stimulation protocol as outlined for the reference AP96 model in section 2.1.2

## 3. Results

In this section, we present how the state space expansion and data-driven models as described in section 2, deal with the two data sets. Since the results for the in-silico synthetic data are easier to interpret, we begin with those data.

### 3.1. Surrogate model for synthetic data

#### 3.1.1. State space

For the data set from numerical simulation of the AP96 model (cf. section 2.1.2), the 3D state space spanned by ***u***= [*u, ũ, g*]^T^ is quite sparse. 95 695 bins in this space contain at least one data point, which corresponds to 9 % of all bins. Of these bins, 40 % contain at least 10 data points. Non-empty bins contain 230 data points on average.

In Fig. 4, we present a three-dimensional view of the state space and two different projections to 2D of the state space; to *u* and *g*, and to *u* and *ũ*, respectively. Panel A shows that most data points lay on a loop-like inertial manifold, while a large volume of the state space is still covered by data points (panel B). Colouring by the mean of 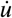, we can see that the two halves of the loop nicely distinguish between excitation and recovery, see the blue and red shades in panel C. Note that the colouring of the state space in panel D corresponds to the SD *σ* and not to the SD *g*. I.e., this panel is coloured by the SD *σ* of the samples of the time-derivative 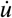 in the bins, instead of the SD *g* of the signal *u* in the neighbourhoods around each point ***x***, which is one of the axes spanning the state space in Fig. 4. In large parts of these projections, the SD *σ* is much lower than the mean absolute value of 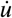, showing that the model’s possible states are well-separated in the chosen state space ***u***.

**Figure 4:**
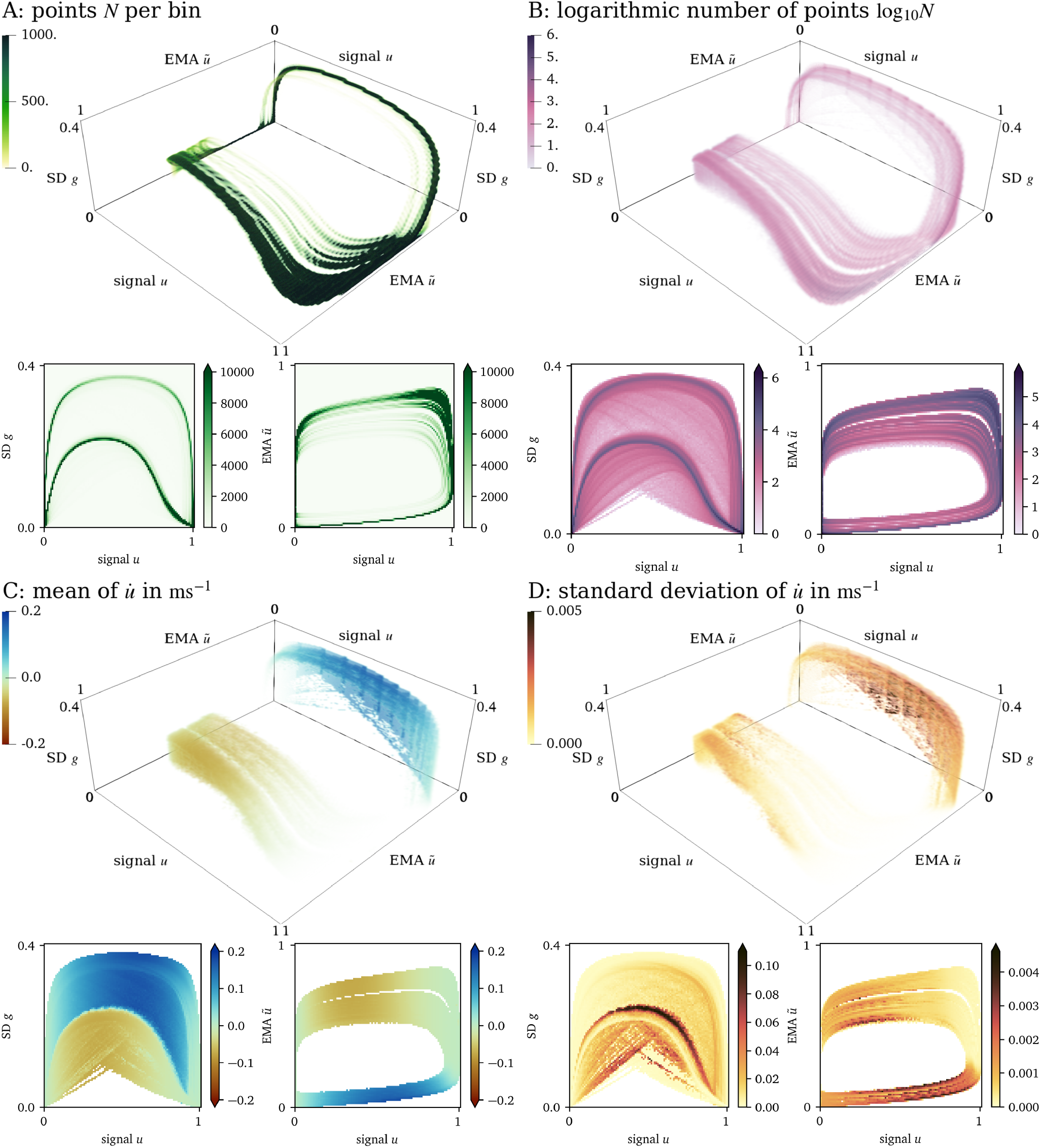
Visualisation of the 3D state space for the AP96 data set and projections onto two dimensions. (A): By colouring the space by the number *N* of points in each bin, it can be seen that most points lay on a loop. (B): Colouring by log_10_ *N*, we can see that still a large volume of the space is covered by data. (C): Looking at the mean of 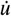 at each bin in this projection, we can see that in one side of the loop, *u* increases while it decreases at the other side. (D): The standard deviation of 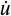 is indicative of whether the states have been chosen well.

#### 3.1.2. Forward model

A model in the context of this work consists of a “colouring” of the entire 3D state space representing the 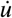 at each possible state ***u***.

The data-driven surrogate model *f* is fit to the 3D state space as outlined in section 2.4 resulting in:

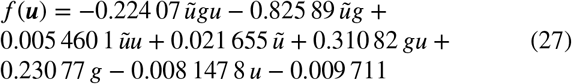

Subsequently, a forward simulation is performed with the model, leading to a prediction of *u* based on the history until that point which can be compared to the reference evolution of *u* in Fig. 5. This simple model displays excitation and recovery, as well as propagation of the wave. The predicted evolution is comparable to the reference: The values for the plateau-phase and resting state are close to the reference values. The depolarisation at the wave front is a bit too late, leading to predicted values that are too low. Likewise, the repolarisation at the wave back is too late, such that there the predicted values are too high.

**Figure 5:**
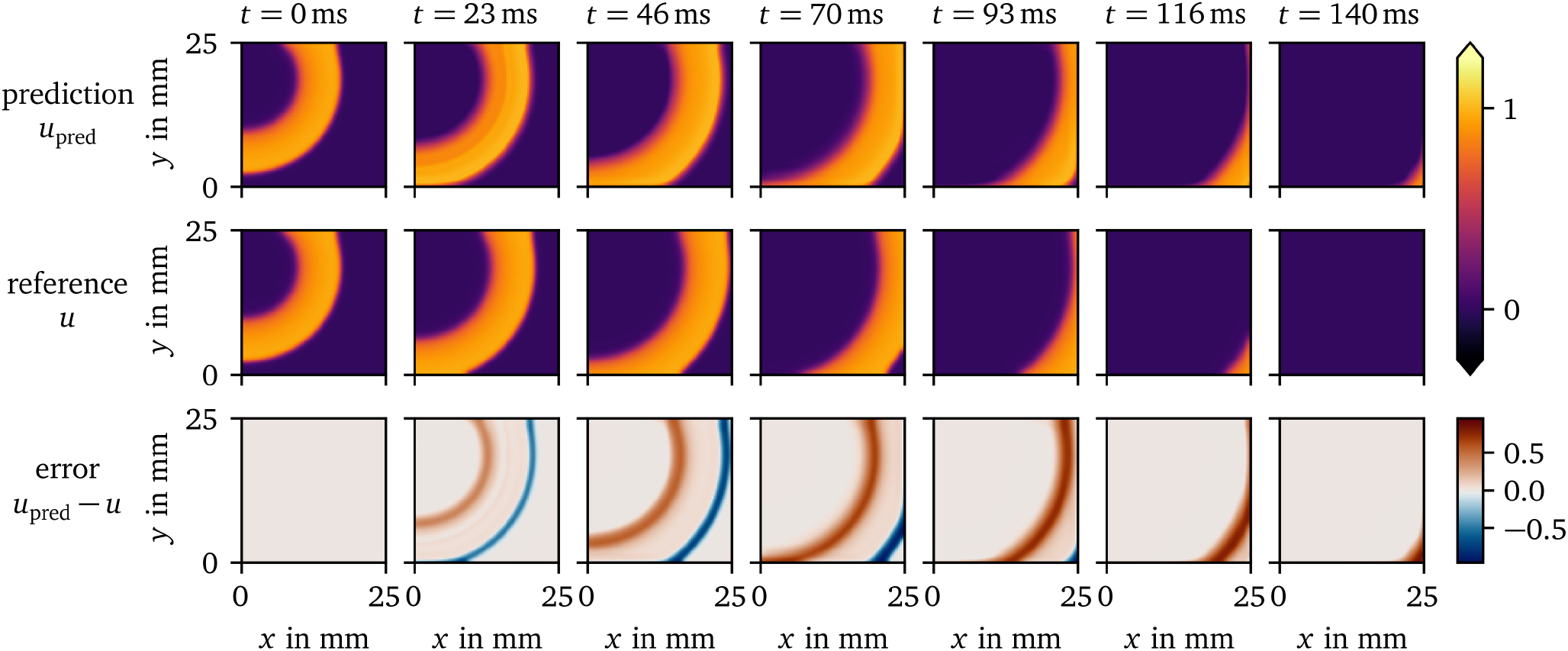
Prediction of the surrogate model over time compared to the reference solution for the AP96 data set. The difference between the prediction and reference is displayed in the bottom row.

Using the method by Bayly et al. [26], conduction velocity has been measured to be CV = (0.1403 ± 0.0036) m s^−1^ for the prediction, and CV = (0.1621 ± 0.0065) m s^−1^ for the reference. This lower propagation velocity leads to the observed delay of wave front and back.

We also computed the average APD of the activity in the central 60 × 60 pixels of the frame with the polarisation threshold of *u* = 0.5: For the prediction, APD = (56.68 ± 0.48) ms can be measured, which is slightly longer than the reference APD = (47.25 ± 0.44) ms.

At the boundary of the domain, the method is still able to compute the evolution well despite less neighbouring data points being used for the calculation of the SD *g*.

#### 3.1.3. Prediction of a spiral wave

We have run simulations with similar initial conditions which lead to spiral waves for the data-driven surrogate AP96 model and its reference. In Fig. 6, the resulting evolutions of *u* over space and time are juxtaposed. It can be seen that the overall wave dynamics are similar, despite the data-driven model only being fit to data from SBP focal waves, i.e., spherical waves growing from a small source region with different timing and amplitudes. The surrogate model predicts wave fronts and backs at similar scales as the reference. The timing of the predicted spiral is also similar to the reference. However, the prediction differs from the reference in the APD of the initial rotation, at the rotor core, and in memory effects. Such a memory effect can clearly be seen in the reference at *t* = 150 ms: The recovery to the resting state is much faster in the rectangle where the second variable *v* of the reference model has been set to a higher value in the initial state of the simulation.

**Figure 6:**
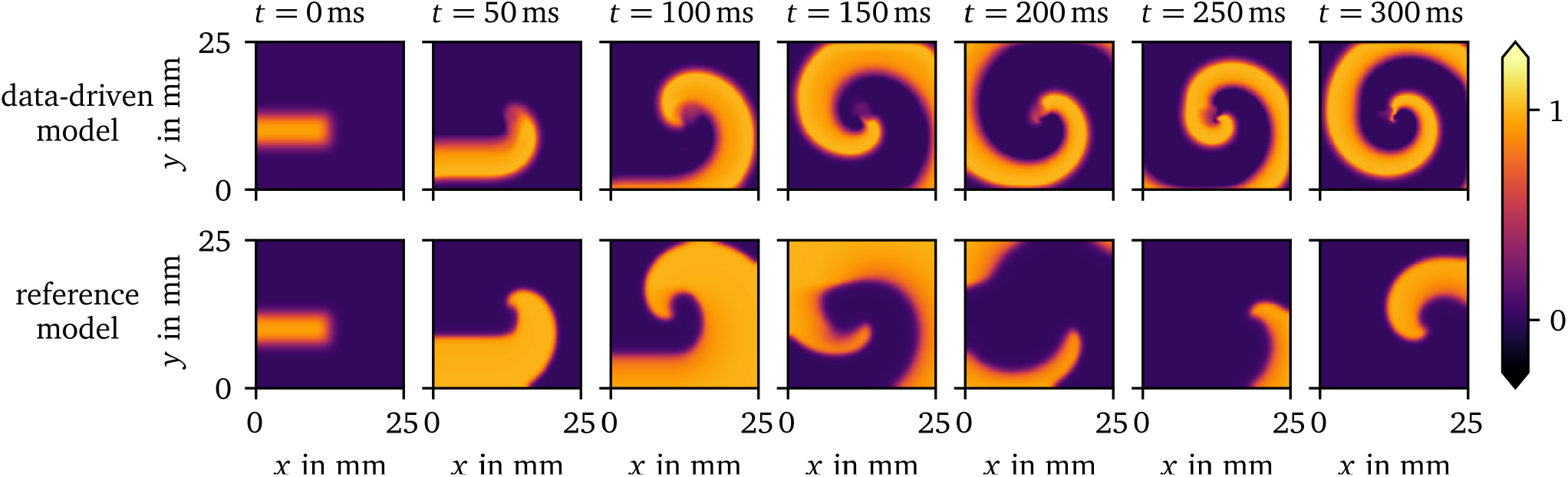
Spiral wave resulting from similar initial conditions for the data-driven surrogate model for the AP96 data set compared to its reference model. The data-driven model can predict the shape of the spiral wave despite only focal waves being included in the training data it is fit to.

#### 3.1.4. Restitution curves

We have also performed 1D cable simulations with the same stimulation protocol for the data-driven surrogate model compared to the rescaled AP96 model for reference. In Fig. 7, the resulting restitution curves of CV (panel A) and APD (panel B) as functions of the CL are shown. The overall shape of the curves for the low-order surrogate model is similar to the reference. However, the predicted CVs are lower than the reference values.

**Figure 7:**
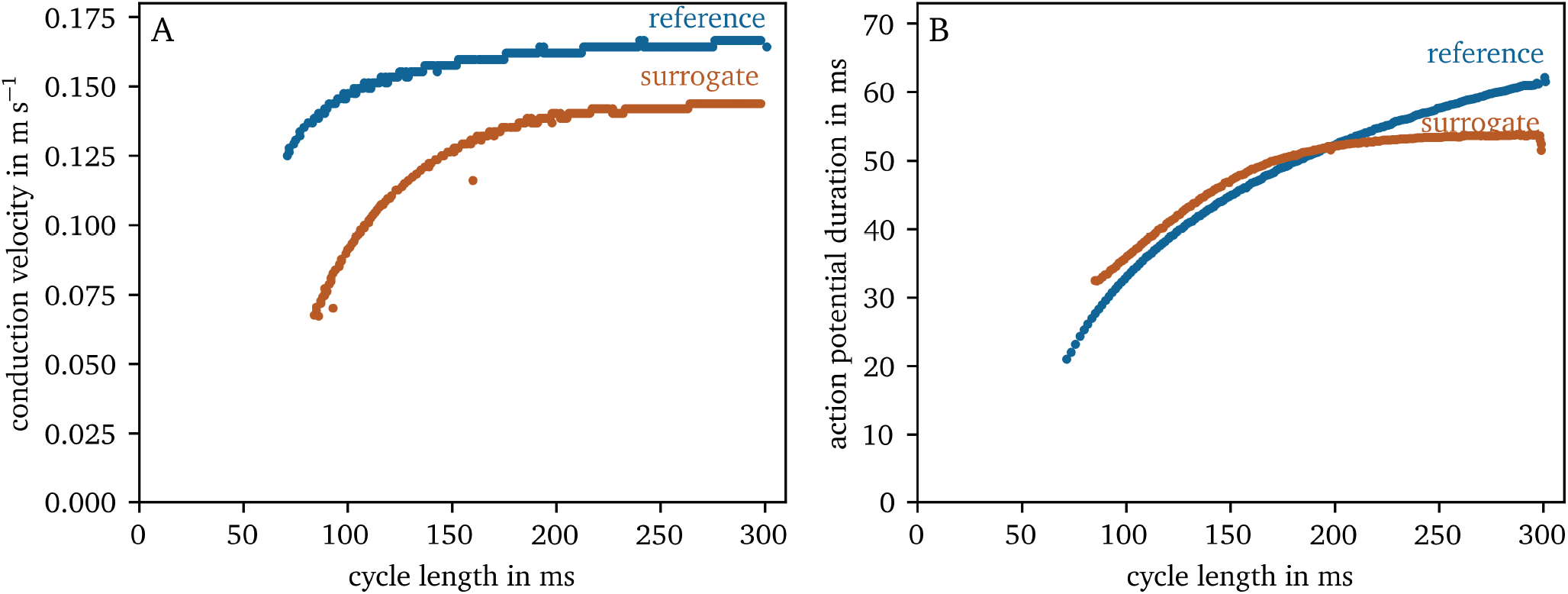
CV and APD restitution curves for the reference AP96 model and its data-driven surrogate model.

### 3.2. Data-driven model for optical mapping data

#### 3.2.1. State space

For the OVM data set of hiAMs (cf. section 2.1.1), the 3D state space spanned by ***u*** = [*u, ũ, g*]^T^ is also sparse: 152 417 bins in this space contain at least one data point, which corresponds to just 15 % of all bins. Of these bins, 71 % contain at least 10 data points. Non-empty bins on average contain 949 data points.

As can be seen in Fig. 8, for these data, we are also observing a loop-like inertial manifold with two halves as in the other data set.

**Figure 8:**
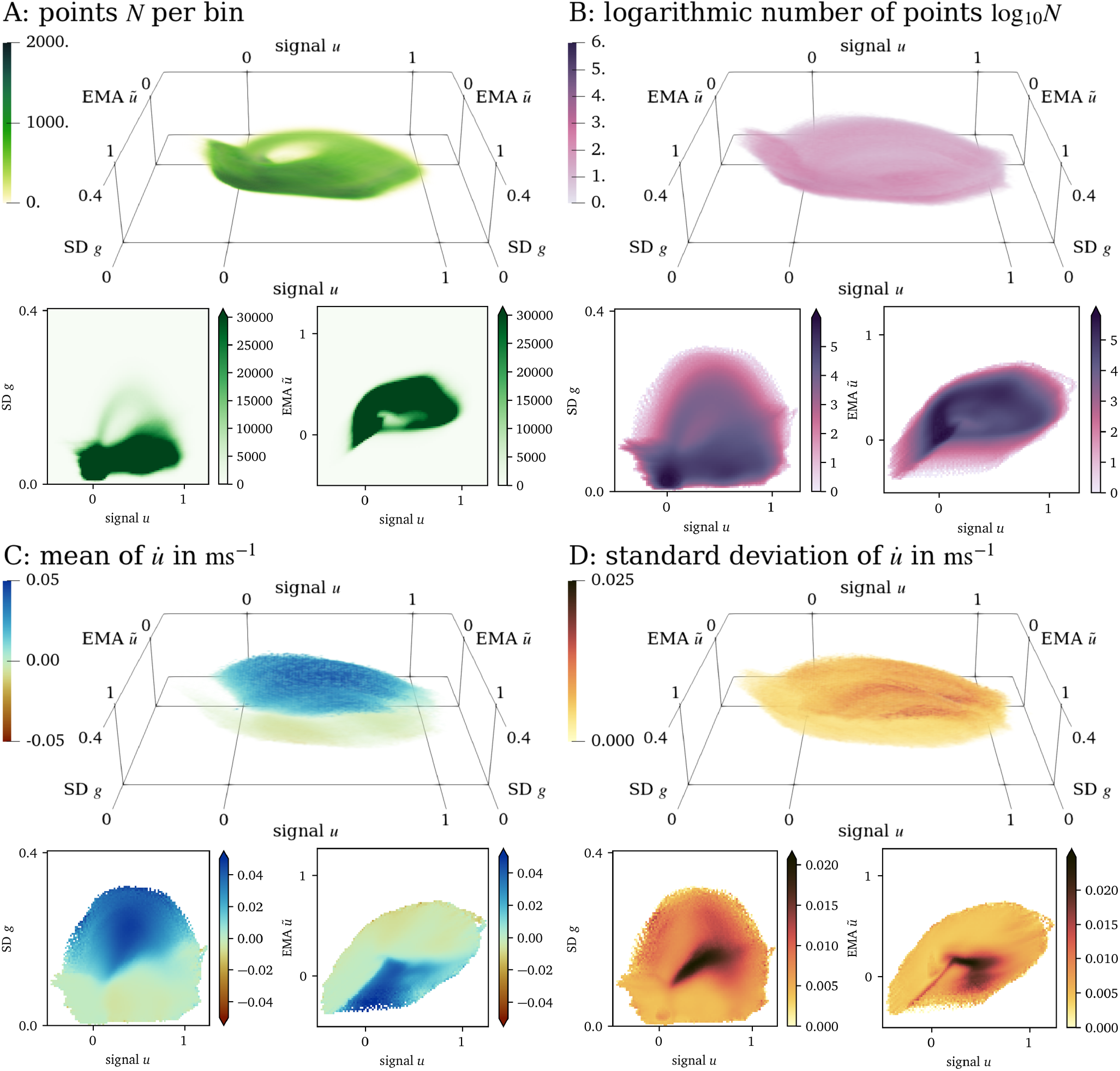
Visualisation of the 3D state space for the OVM data set and projections onto two dimensions in the same style as in Fig. 4.

#### 3.2.2. Forward model

For the OVM data set, the fit polynomial for the data-driven model is:

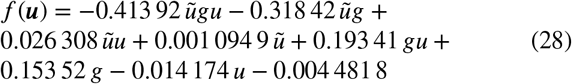

The prediction by the forward model for the OVM data set is depicted in Fig. 9. It models the overall behaviour of the tissue, i.e., excitation, recovery, and wave propagation, at similar time scales: The APD for an excitation threshold of *u* = 0.5 at APD = (140.1 ± 5.1) ms for the prediction is longer than APD = (127 ± 36) ms for the reference. The CV for the prediction is measured at CV = (0.148 ±0.012) m s^−1^, and CV = (0.21 ± 0.13) m s^−1^ for the reference. The large uncertainty in the APD and CV measurements for the reference data is due to small regions of inhomogeneity in the recording. Due to the uncertainty and disparity in APD and CV between prediction and reference, no reliable restitution curves can been measured for this model.

**Figure 9:**
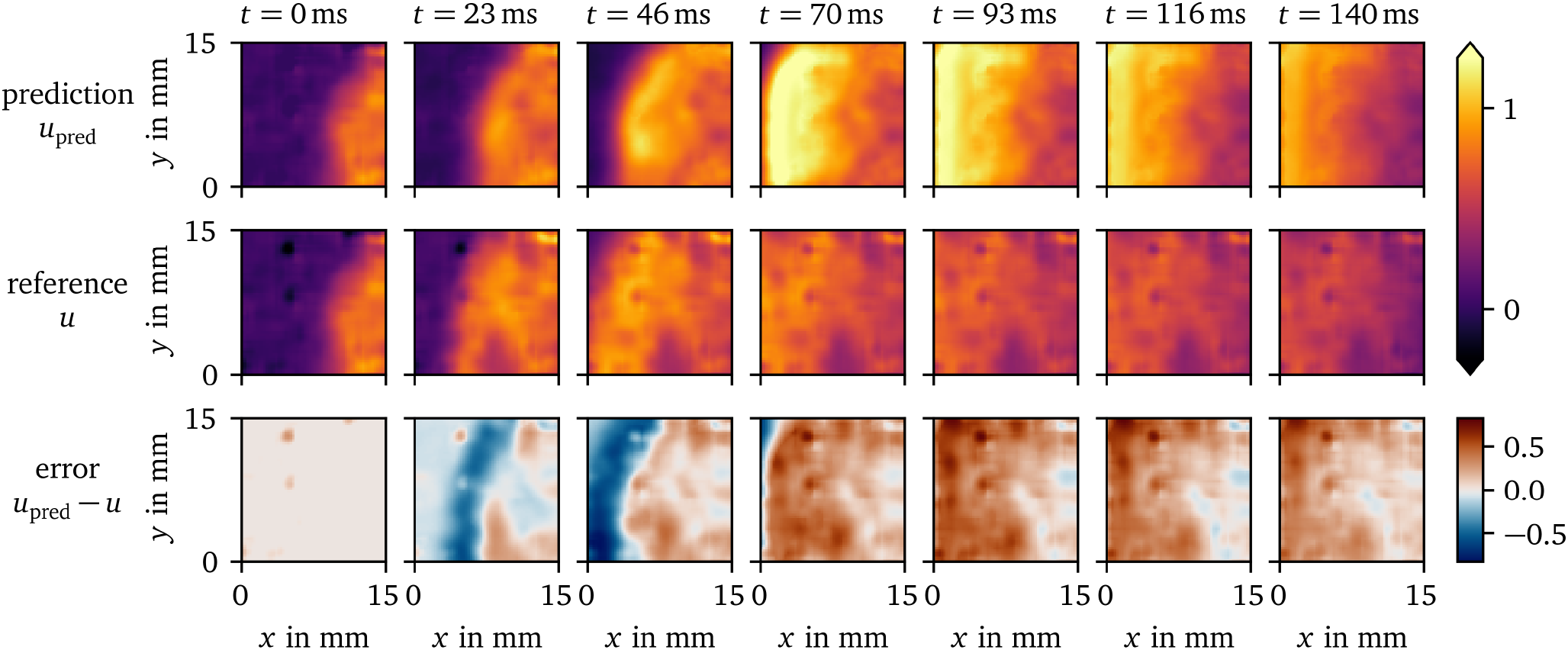
Prediction of the data-driven model over time compared to the reference solution for the OVM data set. The difference between the prediction and reference is displayed in the bottom row.

It can be seen that such artefacts due to inhomogeneity in the monolayer and other more complex effects are difficult to predict for the model which only gets the state *u*(*t*_0,_ *x*) at the initial time as input. Yet, similar structures are found in the prediction.

For the OVM data, the repolarisation at the wave back is much slower than the depolarisation at the wave front. For the synthetic data set AP96 re- and depolarisation happen at roughly the same time scale. Still, the one EMA variable *ũ* is enough to capture these processes in both cases.

#### 3.2.3. Prediction of a spiral wave

In Fig. 10, we compare a rotor in another OVM recording of hiAMs, i.e., additional in-vitro data, and a rotor predicted by the in-silico data-driven model. The model has been fit to data from only focal waves. To stimulate a spiral wave, we use the rectangle-based initial conditions outlined in section 2.1.2. Like for the synthetic data set (AP96, section 3.1), the tissue model is able to predict the overall spiral dynamics. Note that the spiral wave in the OVM recording has stabilised for several minutes before *t* = 0 ms, while the rotor is just forming in the simulation with the data-driven model. The APD predicted by the model is much longer than the observed APD in OVM. This leads to the wave front running into the wave back in the simulation.

**Figure 10:**
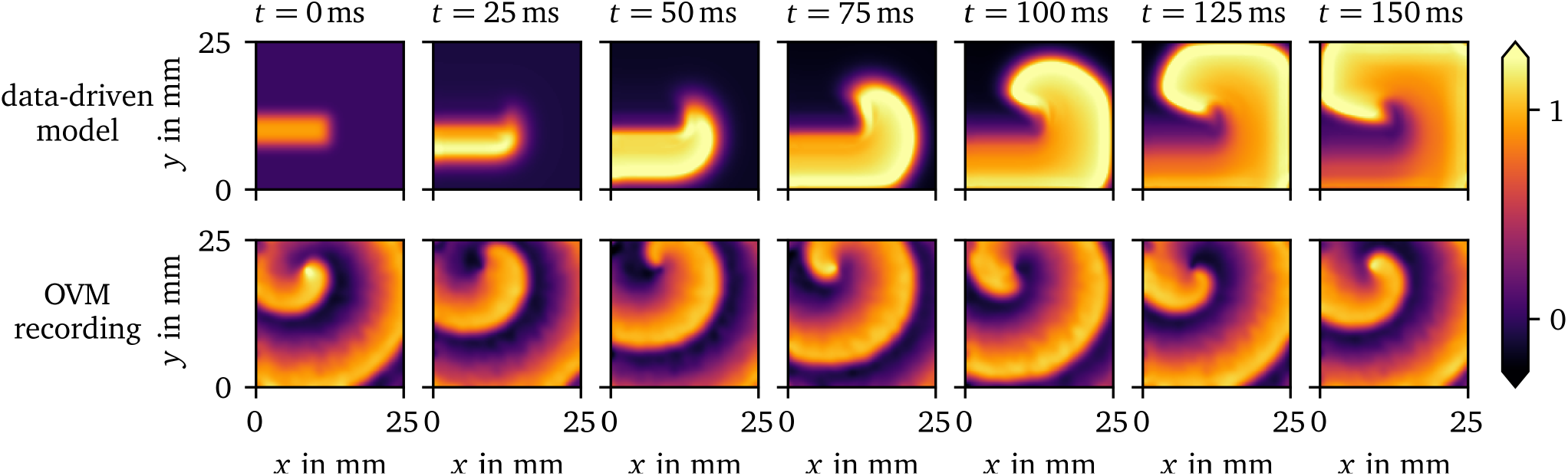
Comparison of a spiral wave for the data-driven model for the OVM data set for hiAMs to a OVM recording of hiAMs showing a stable rotor. The data-driven model can predict the overall dynamics of the spiral wave despite only focal waves being included in the training data it is fit to.

## 4. Discussion

In this paper, we have shown that enough information can be extracted from just one spatio-temporal variable to build a state space, which enables the creation of in-silico models that can predict the further dynamics of the system. In layman’s terms, from a single movie showing the behaviour of excitation and recovery, a predictive model can be constructed. Just the information from OVM is already enough to fit a data-driven model to the dynamics. With our approach, it is even possible to fit a stable model to a single excitation wave, which, in turn, does not generalise well to other waves. For this reason, the SBP protocol is designed to ensure that less likely stimulation patterns are observed in the data, too. The resulting models are able to replicate the excitation patterns of focal waves which are included in the training data and can even predict the dynamics of spiral waves, a pattern that the models are not trained on.

While the variables with which we augment the state space should be seen as preliminary and not optimal choices, the models created with these variables already are able to capture crucial behaviour of the tissue, namely excitation, recovery and wave propagation. Both short and long-term memory can be encoded with the use of EMAs as state variables, which can easily be updated on the fly (eq. 9). It remains to be seen if more or other state space augmentations can be used to be able to replicate higher-order effects, such as wave front curvature and to obtain more accurate restitution behaviour.

The final polynomial model *f* is fit using very a basic supervised machine learning method. However, this is just a small part of the method presented in this work. We have shown that using unsupervised learning, i.e., data transformations, much insight can be gained about the structure of the data. The insights obtained from presenting the data sets in the chosen 3D state spaces as done in this work make it almost trivial to fit the data-driven model to the data.

Through state space expansion with EMAs and SDs, we have found a way to encode useful information to describe the mechanics of excitation waves. It is remarkable, that, with the proper state space expansion, even a simple polynomial is able to capture crucial behaviour of the model sufficiently in the sense that we observe excitation and recovery with an APD similar to the data, and wave propagation at similar CV.

Another useful property of the models created in this work is that they require less computational effort in forward simulations than traditional in-silico models. In the numerical solution of reaction-diffusion systems (eq. 17), a sufficiently small numerical time step must be chosen according to a stability criterion. Typically, this leads to numerical time steps of the order of magnitude 0.1 ms in 2D simulations of in-silico tissue models of cardiac electrophysiology. Forward Euler stepping our model can be done at much larger time steps of up to 10 ms, with the trade-off of fitting less to the original data. Also the computation of the polynomial *f*, EMAs and SDs result in an overall computationally less expensive model than typical tissue models.

While this was not needed for this use case, in further research, more sophisticated general function approximators may be tried for *f*, such as neural networks. Although the Latin hypercube sampling enables some basic regularisation of the solution, it remains to be seen how to keep the fit functions simple enough such that they generalise well. Another limitation that could be addressed when using more flexible function approximators is that, in some cases, the polynomial model *f* updates the state *u* such that it tries to leave the valid range.

While other augmentations were tried in the development of this method, such as pseudo-electrograms, the last APD and diastolic interval at each point over time, or the absolute value of the gradient fit to a neighbourhood of each point using the least-squares method, we are here presenting two augmentations we found most useful, the EMA *ũ* as well as the SD *g*.

We have not yet taken into account boundary effects in the model creation procedure. Some information about the gradient of the main variable *u* is encoded in the SD *g* in the neighbourhood of each point in the medium. The SD *g* can still be computed near the boundary by treating those points just like interior points but with fewer samples. How to encode boundary conditions in this model could be investigated in follow-up studies.

The procedure to generate in-silico tissue models is designed to work well in a setting where not much is known about the wave propagation in an in-vitro tissue model. An advantage of this method is that models for any observed behaviour can be created and fit to data which is in contrast to classical phenomenological models [4–10]. With our method, a data-driven model can be fit with relative ease within minutes that then can be used to predict the further evolution of the tissue. Possible applications include an online-learning approach, where the data-driven model’s predictive power is continuously improved while new data comes in. Such a model could, for instance, be fit to patient recordings, such as intra-cardiac electrograms or electrocardiograms on the body surface, and then be used to predict cardiac arrhythmias before they happen.

## 5. Conclusion

In this work, we presented a method to create simple data-driven in-silico tissue models from in-vitro OVM data and synthetic data that capture the basic behaviour of the corresponding excitable media. These models can predict the activation patterns included in the training data, i.e., only focal waves, and generalise to unseen activation patterns, such as spiral waves. A useful feature of this model creation method is its ability to extract all relevant information that the models are fit to from just one state variable over space and time, such as the transmembrane voltage of cardiomyocytes which can be obtained from a single experiment, e.g. an OVM recording.

## Data availability

Source code of the software implemented for this paper is publicly available at https://gitlab.com/heartkor. The functionality is broken up into several separate packages that may depend on each other.

The model creation pipeline is implemented as the Distephym Python module, `distephym` 1.0.0. It stores the augmented state spaces in sparse histograms, specifically implemented for this publication as another module, `sparsealgos` 1.0.0.

Reading, writing, and analysis of OVM recordings is implemented in the Sappho Python module, `sappho` 1.0.0.

The Ithildin C++ framework, `ithildin` 3.2.2, is a finite-differences, parallelised, reaction-diffusion solver written in C++. It will be released in a separate, dedicated publication [27, 28]. We have used Ithildin to generate the synthetic data set (AP96).

For the other forward simulations in this work, we have written another simple finite-differences reaction-diffusion solver in Python, `ithilmin` 1.0.0, that can easily be combined with other Python modules. It was used for forward runs of the fit data-driven models.

In conjunction with this paper, we are also releasing a new version of the Ithildin Python module, `py_ithildin` 0.6.1, [22]. This module was used for the processing of in-vitro/in-silico data sets. Results from both reaction-diffusion solvers, `ithildin` and `ithilmin`, can be analysed in with this Python module.

The source code of these projects and the AP96 and OVM data sets used in this work have been archived on Zenodo (DOI: 10.5281/zenodo.8183722).

## Acknowledgments

We are grateful to our collaborators at the LUMC, especially to Sven O. Dekker and Juan Zhang for providing access to hiAM monolayers and the OVM setup, and to Niels Harlaar for the spiral wave OVM recording, respectively. Also, we would like to thank all members of Team HeartKOR at KULAK for valuable help during writing, in particular, Marie Cloet.

## CRediT authorship contribution statement

**Desmond Kabus:** Conceptualisation, Methodology, Software, Validation, Formal analysis, Investigation, Data curation, Writing – original draft, Writing – review & editing, Visualisation. **Tim De Coster:** Methodology, Writing – review & editing. **Antoine A.F. de Vries:** Conceptualisation, Resources, Writing – review & editing. **Daniël A. Pijnappels:** Conceptualisation, Resources, Writing – review & editing, Supervision, Project administration, Funding acquisition. **Hans Dierckx:** Conceptualisation, Resources, Writing – review & editing, Supervision, Project administration, Funding acquisition.

## References

[1] Steven A. Niederer, Joost Lumens, and Natalia A. Trayanova. Computational models in cardiology. NATURE REVIEWS CARDIOLOGY, 16(2):100–111, 2019. ISSN 1759-5002. doi: 10.1038/s41569-018-0104-y.

[2] Natalia A. Trayanova, Ashish N. Doshi, and Adityo Prakosa. How personalized heart modeling can help treatment of lethal arrhythmias: A focus on ventricular tachycardia ablation strategies in post-infarction patients. WIREs Systems Biology and Medicine, 12(3), May 2020. ISSN 1939-5094, 1939-005X. doi: 10.1002/wsbm.1477. URL https://onlinelibrary.wiley.com/doi/10.1002/wsbm.1477.

[3] R.H. Clayton, O. Bernus, E.M. Cherry, H. Dierckx, F.H. Fenton, L. Mirabella, A.V. Panfilov, F.B. Sachse, G. Seemann, and H. Zhang. Models of cardiac tissue electrophysiology: Progress, challenges and open questions. Progress in Biophysics and Molecular Biology, 104 (1-3):22–48, jan 2011. doi: 10.1016/j.pbiomolbio.2010.05.008. URL https://doi.org/10.1016%2Fj.pbiomolbio.2010.05.008.

[4] Richard FitzHugh. Impulses and Physiological States in Theoretical Models of Nerve Membrane. Biophysical Journal, 1(6):445–466, July 1961. ISSN 00063495. doi: 10.1016/S0006-3495(61)86902-6. URL https://linkinghub.elsevier.com/retrieve/pii/S0006349561869026.

[5] J. Nagumo, S. Arimoto, and S. Yoshizawa. An Active Pulse Transmission Line Simulating Nerve Axon. Proceedings of the IRE, 50(10): 2061–2070, October 1962. ISSN 0096-8390. doi: 10.1109/JRPROC.1962.288235. URL http://ieeexplore.ieee.org/document/4066548/.

[6] Dwight Barkley. A model for fast computer simulation of waves in excitable media. Physica D: Nonlinear Phenomena, 49(1-2):61–70, April 1991. ISSN 01672789. doi: 10.1016/0167-2789(91)90194-E. URL https://linkinghub.elsevier.com/retrieve/pii/016727899190194E.

[7] Rubin R. Aliev and Alexander V. Panfilov. A simple two-variable model of cardiac excitation. Chaos, Solitons & Fractals, 7(3):293–301, March 1996. ISSN 09600779. doi: 10.1016/0960-0779(95)00089-5. URL https://linkinghub.elsevier.com/retrieve/pii/0960077995000895.

[8] Flavio Fenton and Alain Karma. Vortex dynamics in three-dimensional continuous myocardium with fiber rotation: Filament instability and fibrillation. Chaos: An Interdisciplinary Journal of Nonlinear Science, 8(1):20–47, mar 1998. doi: 10.1063/1.166311. URL https://doi.org/10.1063%2F1.166311.

[9] Alfonso Bueno-Orovio, Elizabeth M. Cherry, and Flavio H. Fenton. Minimal model for human ventricular action potentials in tissue. Journal of Theoretical Biology, 253(3):544–560, aug 2008. doi: 10.1016/j.jtbi.2008.03.029. URL https://doi.org/10.1016%2Fj.jtbi.2008.03.029.

[10] Christopher D. Marcotte and Roman O. Grigoriev. Dynamical mechanism of atrial fibrillation: A topological approach. Chaos: An Interdisciplinary Journal of Nonlinear Science, 27(9):093936, sep 2017. doi: 10.1063/1.5003259. URL https://doi.org/10.1063%2F1.5003259.

[11] G.K. Moe, W.C. Rheinbolt, and J.A. Abildskov. A computer model of atrial fibrillation. Am. Heart J., 67:200–220, 1964.

[12] Marc Courtemanche, Rafael J Ramirez, and Stanley Nattel. Ionic mechanisms underlying human atrial action potential properties: insights from a mathematical model. American Journal of Physiology-Heart and Circulatory Physiology, 275(1):H301–H321, 1998.

[13] Michelangelo Paci, Jari Hyttinen, Katriina Aalto-Setälä, and Stefano Severi. Computational models of ventricular-and atrial-like human induced pluripotent stem cell derived cardiomyocytes. Annals of biomedical engineering, 41(11):2334–2348, 2013.

[14] Rupamanjari Majumder, Wanchana Jangsangthong, Iolanda Feola, Dirk L Ypey, Daniël A Pijnappels, and Alexander V Panfilov. A mathematical model of neonatal rat atrial monolayers with constitutively active acetylcholine-mediated k+ current. PLoS computational biology, 12(6):e1004946, 2016.

[15] Jeffrey J. Fox, Eberhard Bodenschatz, and Robert F. Gilmour. Period-Doubling Instability and Memory in Cardiac Tissue. Physical Review Letters, 89(13):1381 September 2002. ISSN 0031-9007, 1079-7114. doi: 10.1103/PhysRevLett.89.138101. URL https://link.aps.org/doi/10.1103/PhysRevLett.89.138101.

[16] Ning Wei, Yoichiro Mori, and Elena G. Tolkacheva. The role of short term memory and conduction velocity restitution in alternans formation. Journal of Theoretical Biology, 367:21–28, February 2015. ISSN 00225193. doi: 10.1016/j.jtbi.2014.11.014. URL https://linkinghub.elsevier.com/retrieve/pii/S0022519314006663.

[17] Leo Priebe and Dirk J. Beuckelmann. Simulation Study of Cellular Electric Properties in Heart Failure. Circulation Research, 82(11): 1206–1223, June 1998. ISSN 0009-7330, 1524-4571. doi: 10.1161/01.RES.82.11.1206. URL https://www.ahajournals.org/doi/10.1161/01.RES.82.11.1206.

[18] Kirsten HWJ ten Tusscher, Denis Noble, Peter-John Noble, and Alexander V Panfilov. A model for human ventricular tissue. American Journal of Physiology-Heart and Circulatory Physiology, 286(4): H1573–H1589, 2004.

[19] Karli Gillette, Matthias A.F. Gsell, Anton J. Prassl, Elias Karabelas, Ursula Reiter, Gert Reiter, Thomas Grandits, Christian Payer, Darko Štern, Martin Urschler, Jason D. Bayer, Christoph M. Augustin, Aurel Neic, Thomas Pock, Edward J. Vigmond, and Gernot Plank. A framework for the generation of digital twins of cardiac electrophysiology from clinical 12-leads ECGs. Medical Image Analysis, 71: 102080, jul 2021. doi: 10.1016/j.media.2021.102080. URL https://doi.org/10.1016%2Fj.media.2021.102080.

[20] Niels Harlaar, Sven O. Dekker, Juan Zhang, Rebecca R. Snabel, Marieke W. Veldkamp, Arie O. Verkerk, Carla Cofiño Fabres, Verena Schwach, Lente J. S. Lerink, Mathilde R. Rivaud, Aat A. Mulder, Willem E. Corver, Marie José T. H. Goumans, Dobromir Dobrev, Robert J. M. Klautz, Martin J. Schalij, Gert Jan C. Veenstra, Robert Passier, Thomas J. van Brakel, Daniël A. Pijnappels, and Antoine A. F. de Vries. Conditional immortalization of human atrial myocytes for the generation of in vitro models of atrial fibrillation. Nature Biomedical Engineering, 6(4):389–402, jan 2022. doi: 10.1038/s41551-021-00827-5. URL https://doi.org/10.1038%2Fs41551-021-00827-5.

[21] Trine Krogh-Madsen, Peter Schaffer, Anne D. Skriver, Louise Kold Taylor, Brigitte Pelzmann, Bernd Koidl, and Michael R. Guevara. An ionic model for rhythmic activity in small clusters of embryonic chick ventricular cells. American Journal of Physiology-Heart and Circulatory Physiology, 289(1):H398–H413, July 2005. ISSN 0363-6135, 1522-1539. doi: 10.1152/ajpheart.00683.2004. URL https://www.physiology.org/doi/10.1152/ajpheart.00683.2004.

[22] Desmond Kabus, Louise Arno, Lore Leenknegt, Alexander V. Panfilov, and Hans Dierckx. Numerical methods for the detection of phase defect structures in excitable media. PLOS ONE, 17(7):1–31, 07 2022. doi: 10.1371/journal.pone.0271351. URL https://doi.org/10.1371/journal.pone.0271351.

[23] Richard A. Gray, José Jalife, Alexandre V. Panfilov, William T. Baxter, Cándido Cabo, Jorge M. Davidenko, and Arkady M. Pertsov. Mechanisms of Cardiac Fibrillation. Science, 270(5239):1222–1223, November 1995. ISSN 0036-8075, 1095-9203. doi: 10.1126/science.270.5239.1222. URL https://www.science.org/doi/10.1126/science.270.5239.1222.

[24] Pawel Kuklik, Stef Zeemering, Bart Maesen, Jos Maessen, Harry J. Crijns, Sander Verheule, Anand N. Ganesan, and Ulrich Schotten. Reconstruction of Instantaneous Phase of Unipolar Atrial Contact Electrogram Using a Concept of Sinusoidal Recomposition and Hilbert Transform. IEEE Transactions on Biomedical Engineering, 62(1):296–302, January 2015. ISSN 0018-9294, 1558-2531. doi: 10.1109/TBME.2014.2350029. URL http://ieeexplore.ieee.org/document/6880777/.

[25] Michael D McKay, Richard J Beckman, and William J Conover. A comparison of three methods for selecting values of input variables in the analysis of output from a computer code. Technometrics, 42(1): 55–61, 2000. Publisher: Taylor & Francis.

[26] Philip V Bayly, Bruce H KenKnight, Jack M Rogers, Russel E Hillsley, Raymond E Ideker, and William M Smith. Estimation of conduction velocity vector fields from epicardial mapping data. IEEE transactions on biomedical engineering, 45(5):563–571, 1998.

[27] Marie Cloet, Louise Arno, Desmond Kabus, Joeri Van der Veken, Alexander V. Panfilov, and Hans Dierckx. Scroll Waves and Filaments in excitable Media of higher spatial Dimension. 2023. doi: 10.48550/ARXIV.2304.14861. URL https://arxiv.org/abs/2304.14861. Publisher: narXiv Version Number: 1.

[28] Desmond Kabus, Marie Cloet, Christian Zemlin, and Hans Dierckx. The ithildin anisotropic reaction-diffusion solver for excitable media. 2023. forthcoming.

